# Training the next generation of researchers in the Organ-on-Chip field

**DOI:** 10.1101/2022.08.03.502617

**Authors:** Alessia Moruzzi, Tanvi Shroff, Silke Keller, Peter Loskill, Madalena Cipriano

## Abstract

Organ-on-chip (OoC) technology bridges the principles of biology and engineering to create a new generation of *in vitro* models and involves highly interdisciplinary collaboration across STEM disciplines. Training the next generation of scientists, technicians and policy makers is a challenge that requires a tailored effort. To promote the qualification, usability, uptake and long-term development of OoC technology, we designed a questionnaire to evaluate the key aspects for training, identify the major stakeholders to be trained, their professional level and specific skillset. The 151 respondents unanimously agreed on the need to train the next generation of OoC researchers and that the training should be provided early, in interdisciplinary subjects and throughout the researchers’ career. We identified two key training priorities: (i) training scientists with a biology background in microfabrication and microfluidics principles and (ii) training OoC developers in pharmacology/toxicology. This makes training in OoC a transdisciplinary challenge rather than an interdisciplinary one. The data acquired and analyzed here serves to guide training initiatives for preparing competent and transdisciplinary researchers, capable of assuring the successful development and application of OoC technologies in academic research, pharmaceutical/chemical/cosmetic industries, personalized medicine and clinical trials on chip.

## Introduction

The last decade has seen the emergence of Organ-on-Chip (OoC) systems, which are defined as “fit-for-purpose microfluidic devices, containing living engineered organ substructures in a controlled microenvironment, that recapitulate one or more aspects of the organ’s dynamics, functionality and (patho)physiological response in vivo under real-time monitoring” (Mastrangeli, Millet, & van den Eijnden-van Raaij, 2019). The technology promises great potential for human-based translational (Rudmann, 2019) and personalized medicine (Van Den Berg et al., 2019) for the reduction of pharmaceutical R&D costs (Franzen et al., 2019; Peck et al., 2020) and for the reduction and replacement of animal models (3Rs). The OoC field is characterized by a high degree of interdisciplinarity and draws upon technologies and concepts from various fields. Approaches from engineering, physics and chemistry enable precise spatiotemporal manipulation and monitoring of cells and liquids. Techniques from molecular and cell biology allow for the recapitulation and investigation of cell and tissue function as well as accurate cellular microenvironments (Sohn et al., 2020). Guidelines from clinical science, toxicology and pharmacology define the applications of these models (OECD, 2018). The emergence and development of the OoC field stems from significant progress in related fields: Advances in microfabrication techniques enabled the generation of polymer structures on cellular scales. Breakthroughs in stem cell research paved the way for fully human tissue models. New approaches in molecular biology and imaging allowed for small-scale and high-throughput readouts. The development of novel sensor technologies enabled long-term monitoring of living systems. OoC technology allows for the generation of mechanistic knowledge that can be a game-changer towards the 3Rs and the transition to non-animal science (Herrmann et al., 2019; Hippenstiel et al., 2021).

The OoC field, however, is still in its infancy, with a variety of challenges impeding the broad adoption of OoC models in academic, clinical and industrial research (Baran et al., 2022; Marx, 2020; Mastrangeli, Millet, Mummery, et al., 2019). Challenges such as the recapitulation of complex tissues, cell maturity, large scale chip production, quality assurance, standardization, facilitation of usability, regulatory acceptance and many others are yet to be addressed (Allwardt et al., 2020; Baran et al., 2022; Mastrangeli, Millet, Mummery, et al., 2019; Mastrangeli, Millet, & van den Eijnden-van Raaij, 2019; Mastrangeli & van den Eijnden-van Raaij, 2021; Peck et al., 2020; Piergiovanni et al., 2021).

To overcome these challenges, it is imperative to train experts in this multidisciplinary technology at different levels of education and work function. Importantly, one must consider that experts have different STEM identities (Singer et al., 2020), distinct primary STEM education fields that can range across physics, biology to engineering, different ways to approach problems and different perceptions depending on their experience level (Stella et al., 2019). Thus, targeted and well-defined training strategies that foster interdisciplinary work and communication for professionals must be defined and implemented. Also, early training initiatives targeting undergraduate and graduate students should aim at integrating disciplines in an transdisciplinary rather than an interdisciplinary context (Takeuchi et al., 2020).

The assessment of training needs has been performed previously in emerging fields such as data analysis (Federer et al., 2016), metabolomics (Weber et al., 2015) and applied health research (Barratt & Fulop, 2016). Training efforts in novel technologies must consider a combination of theoretical and practical immersive pedagogy with exposure to workflows and processes and addition to curricula. Moreover, such training efforts should also encompass non-technical topics such as bioethics, data privacy, trial management and personal safety (Sánchez Carracedo et al., 2018).

The aim of this study is the identification of the training priorities, key skill sets and adequate types of training required to prepare the next generation of researchers to advance the OoC field and possibly change the way we perform biomedical, pharmaceutical and toxicological research. Researchers working in the OoC field take on many different roles: *scientists as developers, scientists as end-users* in *academia* and *industry, technicians, clinicians*, and *scientists as decision makers*. Each of these “stakeholders” requires a specific amount and type of training to successfully adopt OoC technology. To identify stakeholder-specific requirements, we conducted a survey aimed at providing an objective assessment of training needs. Experts actively working in the OoC field from different sectors, disciplines and career stages were asked to complete a questionnaire focused on three main overarching questions:

- Which **aspects of training** are most important for the overall development of the OoC field?
- Which **specific type of training** in the OoC field should each stakeholder receive?
- What is the **amount of training** that each stakeholder should receive to achieve a certain level of experience in the OoC field?

## Methods

### Survey description and recruiting method

The evaluation of training needs was based on a structured questionnaire accessible via the *EUSurvey* website from 01/15/2019 to 06/12/2020 (ec.europa.eu/eusurvey/). Stakeholders from the OoC field were invited to participate via direct email, newsletters and social media (*e.g*., Twitter, LinkedIn, ResearchGate). The questionnaire (**supplementary file 1**) comprised of 15 questions and was completed by 151 respondents. 122 respondents answered all questions. **Supplementary file 2** includes a table with an overview of the number of respondents that answered each question, as well as all analyzed data used to generate the figures. The specific questions in the survey are listed below:

- Q1. How would you define yourself as a professional? a) Type of institution; b) Job Level; c) Field of work; d) Field of Graduate University education (Bachelor’s Degree); e) Field of first Postgraduate University education (Master’s Degree); f) Field of second Postgraduate University education (Doctorate Degree); g)Main area of expertise
- Q2. Select the tissues/organs/systems/biological functions with which you are familiar and/or you work with? Q3. Select the microfabrication techniques for polymer-based microfluidic devices with which you are familiar and/or you work with? Q4. How important are the following aspects for the Organ-on-Chip field development? Q5. How important is it to provide specific training for each of the following stakeholders, to promote the Organ-on-Chip systems qualification, usability, uptake and/or long-term development? Q6. At which level do you consider that specific training is necessary to promote the Organ-on-Chip systems qualification, usability, uptake and/or long-term development? Q7-12. How important are the following elements to consider for training, for Scientists as developers/end users (Academia/Industry)/decision makers/Technicians/Clinicians regarding the improvement of Organ-on-Chip systems qualification, usability, uptake and/or long-term development? Q13. What is the most adequate complexity level of specific training for each of the following stakeholders, to promote Organ-on-Chip systems qualification, usability, uptake and/or long-term development? Q14. What is the most adequate amount of specific training for each of the following stakeholders, to promote Organ-on-Chip systems qualification, usability, uptake and/or long-term development?
- Q15. How important would it be to include the topic of Organ-on-Chip technologies as a seminar of course in the following broader field(s) of education?

### Methods of data analysis

The raw data was exported from the *EUSurvey* platform as a Microsoft Excel spreadsheet and was further analyzed in Excel 2019 using power query scripts to perform comparative analyses. The raw data is included in **supplementary file 3** with the integrated data analysis scripts.

Given the breadth of survey responses, certain factors were taken into consideration during data analysis. First, answers from respondents who did not specify their *Job level* or selected multiple Job levels were excluded from the analysis, thus reducing the number of respondents from 151 to 149. Second, whenever a respondent selected the option “Other (please specify)____” and typed in the specific answer, we assessed the response on a case-by-case basis and determined whether to included it in one of the available options or to categorize it indeed as *other*. This is described in detail in the “Substitution Tables” (**supplementary file 3)**.

Q1, Q2, Q3, Q6, Q14 and Q15 were analyzed by counting the number of respondents for each option. Because multiple answers were allowed, the total number of answers outnumbers the total number of respondents. Additionally, we counted how many respondents from Q2 and Q3 were familiar with more than one Microfabrication technique or more than one organ.

For Q6, we also analyzed which combinations of educational levels (Undergraduate, Master, Doctorate and Postdoctoral) were selected by most respondents.

The questions asking to grade the degree of importance (Q4, Q5, Q7-12 and Q15) were analyzed by assigning values to the options to generate a ranking as follows: Very important =4; Somewhat important = 3; Less important = 2; Not important = 1. This ranking system was chosen to reveal the highest contrast across responses within the different categories, thus enabling clear conclusions to be drawn from the graphs. We excluded respondents who skipped a question and those who answered “not sure”. Q15 was available only to those respondents who considered it important to include training in a specific course for at least one stakeholder in Q14. The data was plotted as heatmap of the ranking number and wherever relevant, the proportion of respondents was included in the text.

### Subgrouping of the respondents

The respondents were categorized in accordance with:

- Work experience: ***PhD candidates*** 46% (69 respondents) **versus *senior scientists*** 51% (79 respondents) – respondents pursuing a doctoral degree compared to those of senior scientists (*e.g*., postdoctoral researchers, department heads, professors and upper management).
- Type of employer: ***Scientists in academia* versus *scientists in non-academic institutions (NAI)*** (regardless of whether they were PhD candidates or Seniors scientists) - respondents from academic institutions compared to those working in industry, hospitals, small and medium-sized enterprises (SMEs, as defined by H2020) governmental and non-governmental organizations. As multiple answers were possible, whenever a respondent selected “academia”, he/she was excluded from the “NAI” group, resulting in 113 respondents in the “academia” group and 38 in the “NAI” group.
- Main area of expertise: “**experts with biology expertise**” (66 respondents that selected <u>only</u> choices from the biology-based area), “**experts with engineering expertise**” (43 respondents that selected <u>only</u> choices from the engineering-based area) or “**experts with interdisciplinary expertise**” (38 respondents that selected at least one biology-based <u>and</u> one engineering-based area of expertise),

These categories were then analyzed with the same power query code written for the analysis of the overall data and compared to each other.

### Graphical representation of the data

GraphPad Prism version 9.3.1 (471) for Windows (64-bit), GraphPad Software, San Diego, California USA (www.graphpad.com) was used solely to generate the graphical representation of the data. No statistical analysis was performed.

## Results

### Professional profile of the respondents

The answers to Q.1 of the survey revealed a diverse group of respondents with respect to their level of profession, their employer type and their main areas of expertise. Forty six percent of the respondents were *PhD candidates* and 51% comprised of *senior scientists* (**Fig.1 a**).

**Figure 1:**
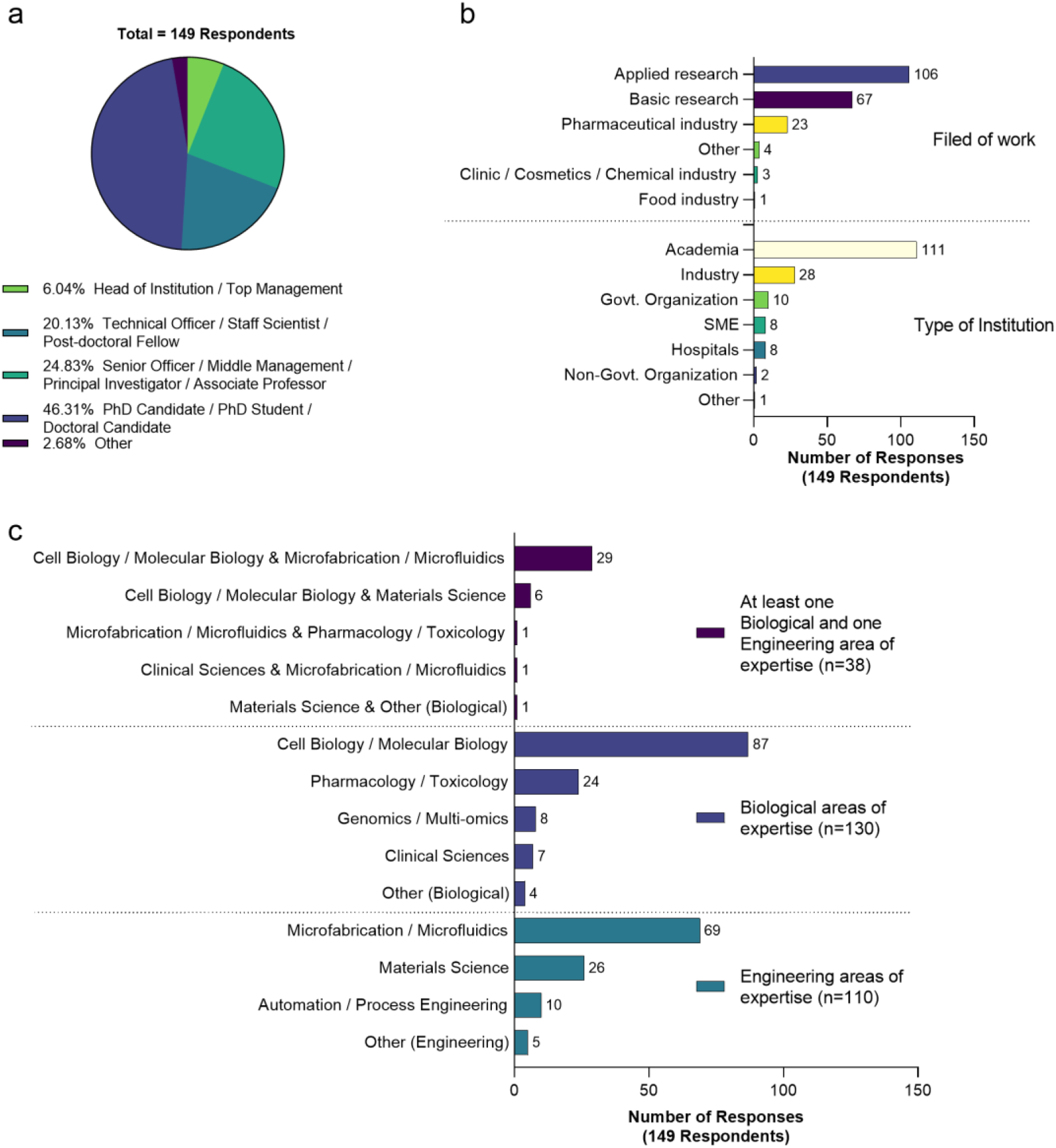
Professional profile of the respondents: (a) Current professional level (b) Current type of employer (c) Areas of expertise, categorized by biology/engineering/interdisciplinary skills. Data in (b) and (c) is from questions in the questionnaire that allowed for multiple answers. The number of respondents is reflected in the figure legend and x axis while the number of responses is described near each bar. The data reflects the responses to Q1.b (a), Q1.a (b) and Q1.c (B), as well as Q1.g (c)

With respect to the current employer, the most represented category was *Academia* (75%), which comprised of universities and research institutes. Amongst *NAI, industry* was the most represented (19%) followed by *governmental organizations* (7%), *SMEs* (5%) and *hospitals* (5%) (**Fig.1 b**). Also, most respondents worked in applied research, as expected in a multidisciplinary field. The main areas of expertise in biology and engineering were *cell and molecular biology* (58%) and *microfabrication//microfluidics* (46%), respectively. A quarter of the respondents selected at least one engineering and one biological option as main areas of expertise (**Fig.1 c**).

### Opinion of the respondents on the state of Organ-on-chip development

Next, we explored which aspects of OoC technology the respondents deemed to be important for the development of the OoC field (**Fig.2**). These aspects were related to design or production (*e.g., usability* or *production scale-up*) and to the translation of OoC technology (*e.g., comparison with clinical data* or *qualification of models*). Undeniably, many of the respondents considered *training* to be very important (100/149) or somewhat important (41/149), highlighting the importance of this study. The top three aspects considered very important, apart from *training* itself, were *definition of cell culture standards – function and origin of cells (112/149), usability* (109/149) and *qualification of the models* (95/149).

**Figure 2:**
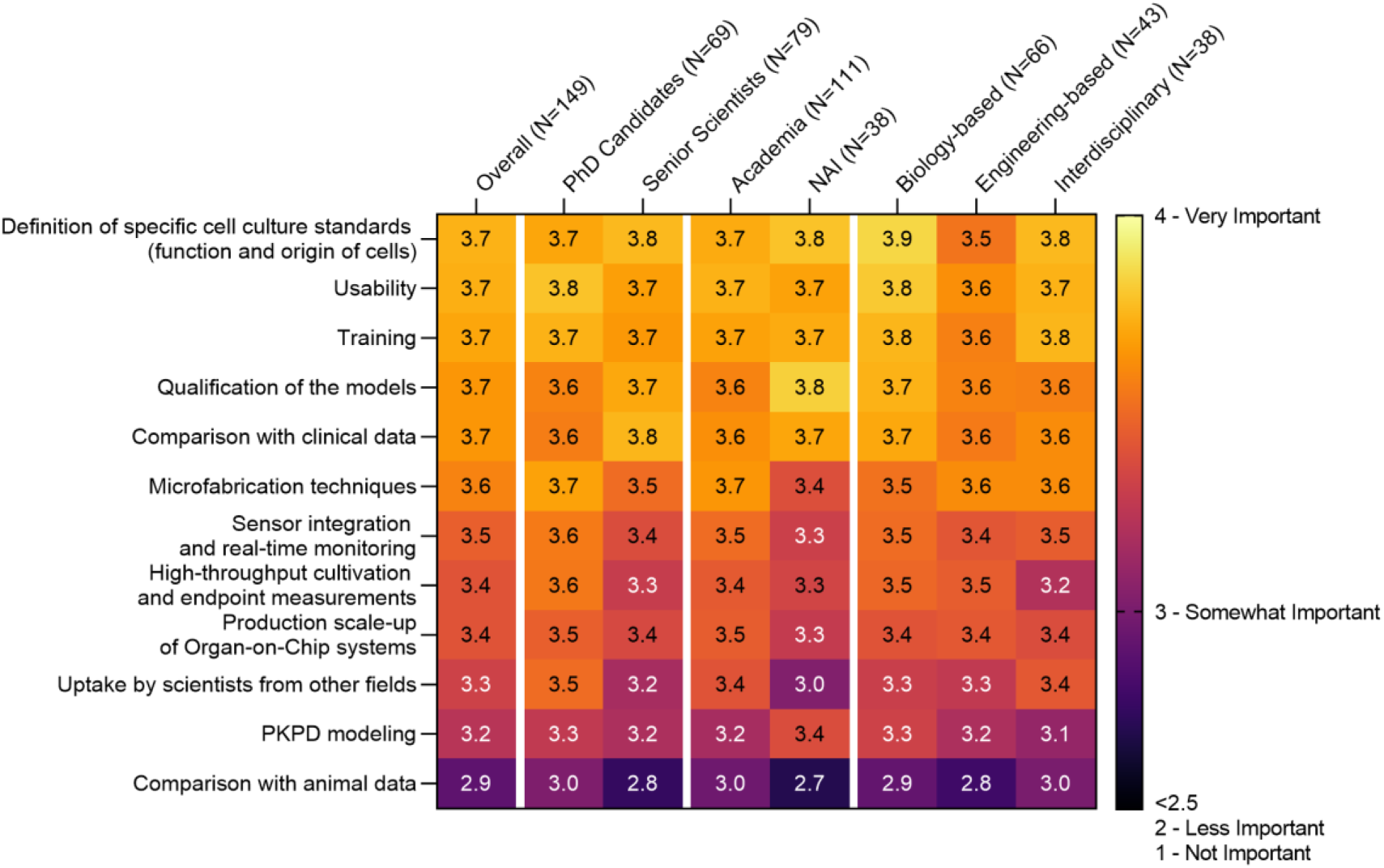
Assessment of the importance of aspects in OoC field development, from left to right for all respondents (overall dataset, N=149), for the subgroups PhD candidates vs. Senior Scientists (N=69+79), for the subgroups Academia vs. NAI (N=111+38), and categorized based on main expertise biology/engineering/interdisciplinary experts (N=66+43+38). N= number of respondents, total and for each subgroup. The importance level was ranked by assigning values to the options to generate a ranking as following: Very important = 4; Somewhat important = 3; Less important = 2; Not important = 1. We excluded respondents who skipped the question and those who answered “not sure”. The heatmap contains the average ranking per response per group. The data reflects the responses to Q4.

PhD candidates considered usability (53/69) as the most important aspect in contrast to senior scientists, who chose definition of cell culture standards – function and origin of cells (63/79) and comparison to clinical data (35/79). Scientists in academia considered usability very important (82/111) while NAI respondents deemed the definition of cell culture standards – function and origin of cells (31/38) as the most important aspects, followed by qualification of the models (29/38). Biological experts, like interdisciplinary experts, considered definition of cell culture standards – function and origin of cells as the most important aspect, followed by usability (52/66 and 29/38, respectively). Engineering experts concurrently identified microfabrication techniques, usability and qualification as the most important aspects (27/43).

Stakeholder groups perform different roles and utilize OoC technology differently. Hence, they require different types of training, as reflected by the responses (**Fig.3**). As expected, the overall responses showed that the key aspects (rank > 3.5/5) for the training of *scientists as developers* were mostly engineering aspects (*biomaterials and microfluidic principles*) and *cell culture and stem cell technology*. For *scientists as end-users in academia* the key aspects were *monitoring and analyzing (sensors, imaging, molecular biology/omics)*, while for *scientists as end-users in industry, pharmacology and toxicology principles*, and *quality assurance* were at the forefront. Translational aspects such as *regulatory affairs, quality assurance* and *ethics* were the most important training aspects for *scientists as decision makers (rank of >3.5/5)*, while the training for *technicians* could focus on *cell culture and stem cell technology*. The respondents also suggested that *clinicians* should be trained in *pharmacology and toxicity principles* and *ethics*, ranking most of the other aspects as less important (rank <3.4/5). Interestingly, *PhD candidates* identified a need for training in *monitoring and analyzing (sensors, imaging, molecular biology/omics), pharmacology and toxicology principles and quality assurance (rank of 3.7/5)* as a priority for *scientists as end-users in industry, conversely to senior scientists* that identified *pharmacology and toxicity principles (rank of 3.6/5)*. Overall, higher importance ranks were given by respondents with *interdisciplinary* expertise compared to *biology* or *engineering* experts, especially for topics such as *monitoring and analyzing (sensors, imaging, molecular biology/omics), pharmacokinetic/pharmacodynamic (PKPD) modeling, pharmacology and toxicology principles* and *quality assurance*. The opinion of respondents working in *NAI* also differed from *Academia* by considering training in *microfabrication* for *scientists as developers* less important but attributing a higher rank to *cell culture and stem cell technology*.

**Figure 3:**
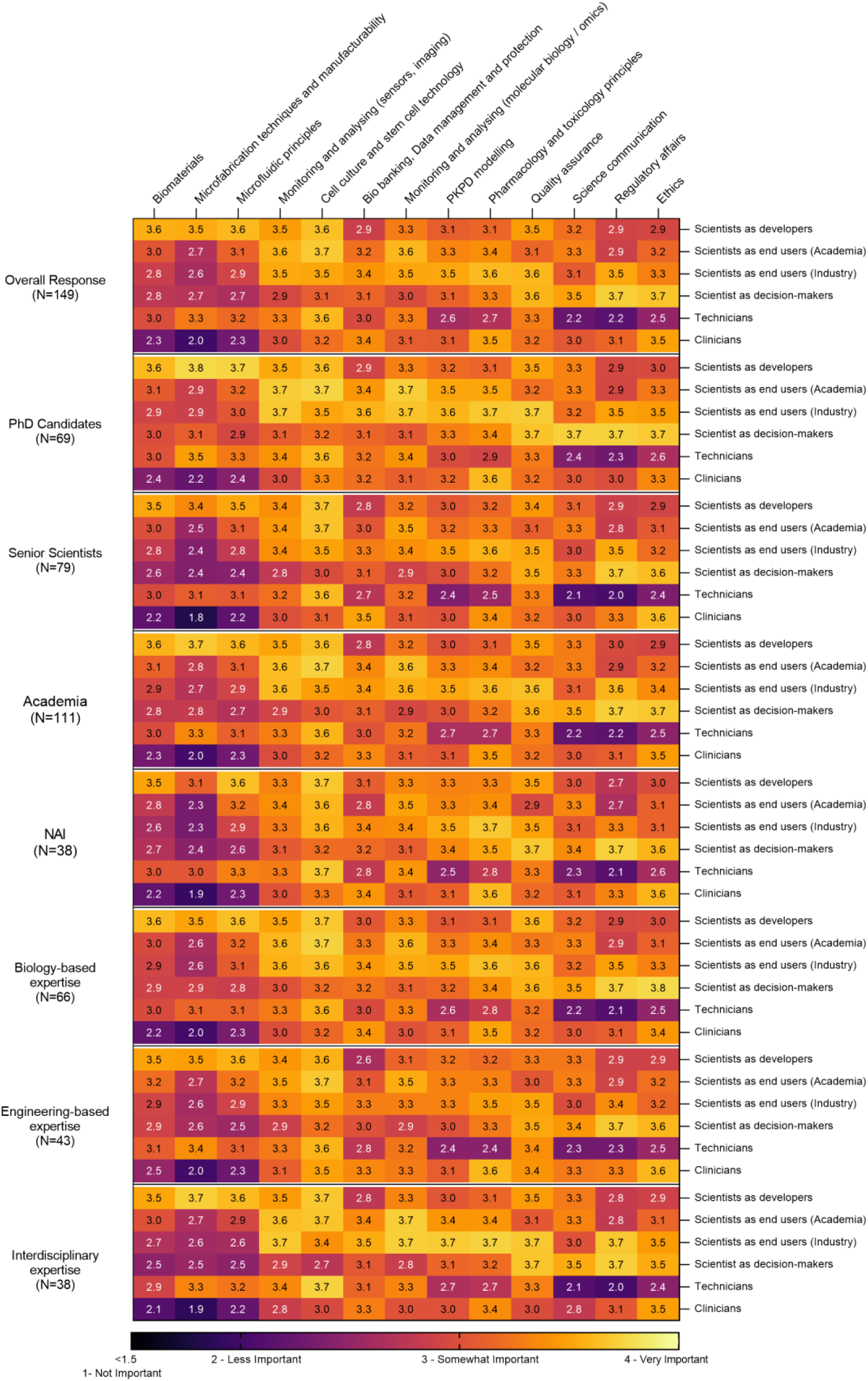
Aspects of training in OoC technology to be provided for the various stakeholders, opinions of all respondents (overall dataset, N=149), PhD candidates vs. senior scientists (N=69+79), biology/engineering/interdisciplinary experts (N=66+43+38) and Academia vs. non-academic institutions (NAI) (N=111+38). The importance level was ranked by assigning values to the options to generate a ranking as following: Very important = 5; Somewhat important = 4; Less important = 3; Not important = 1. We excluded respondents who skipped the question and those who answered “not sure”. The heatmap contains the average ranking per response and per group. The data reflects the responses to Q7-12. Engineering aspects include Biomaterials, microfabrication techniques and manufacturability, microfluidic principles, and monitoring and analysing (sensors, imaging). Biological aspects include cell culture and stem cell technology, biobanking, data management and protection, monitoring and analysing (molecular biology/omics), PKPD modelling, and pharmacology and toxicology principles. Translational aspects include quality assurance, science communication, regulatory affairs and ethics.

### Respondents Opinion on specific training needs

The next question in the survey addressed the nature of the training to be provided to each stakeholder. Respondents were asked at which stage of their education the stakeholders should receive training, the duration of the training itself and in which fields of education OoC-specific training should be integrated.

Overall, training was deemed necessary in all stages of education (**Fig.4 a)**, particularly at a *master’s* and *doctorate level* (70% and 81% respectively). Only 1% of the respondents believed that training should begin only at a *postdoctoral level*, suggesting focus on OoC training at early stages of education. The respondents ranked as very important, the integration of OoC training in *bioengineering* degrees (122/144), followed by *pharmacology/toxicology (86/120), while chemistry (13/119)* was considered a low priority (**Fig.4 b**). With respect to the intensity of the training, there was a strong agreement (79 %) that *scientists as developers* should be trained in a specific *postgraduate course (1 to 2 years)* to develop deep knowledge and acquire practical skills (**Fig.4 c**). *One semester of training* was recommended by respondents for *scientists as end-users in academia* (59%) and for *scientists as end-users in industry* (45%). However, 45% of the respondents considered *20 hours of practical training* adequate for *scientists as end-users in industry*. For *scientists as decision makers*, 48% of the respondents recommended gaining theoretical competence through *20 hours of non-practical training*. For *technicians, 20 hours of practical training* were deemed enough to achieve competence in practical skills by most of the respondents (66%). With respect to *clinicians*, there was a consensus that they only require an *introductory awareness by receiving up to 20 hours of training* but the difference lays in the nature of training being *practical* (35%) or *non-practical training* (45%). The recommendation for the length of training programs progressively increased with the degree.

**Figure 4:**
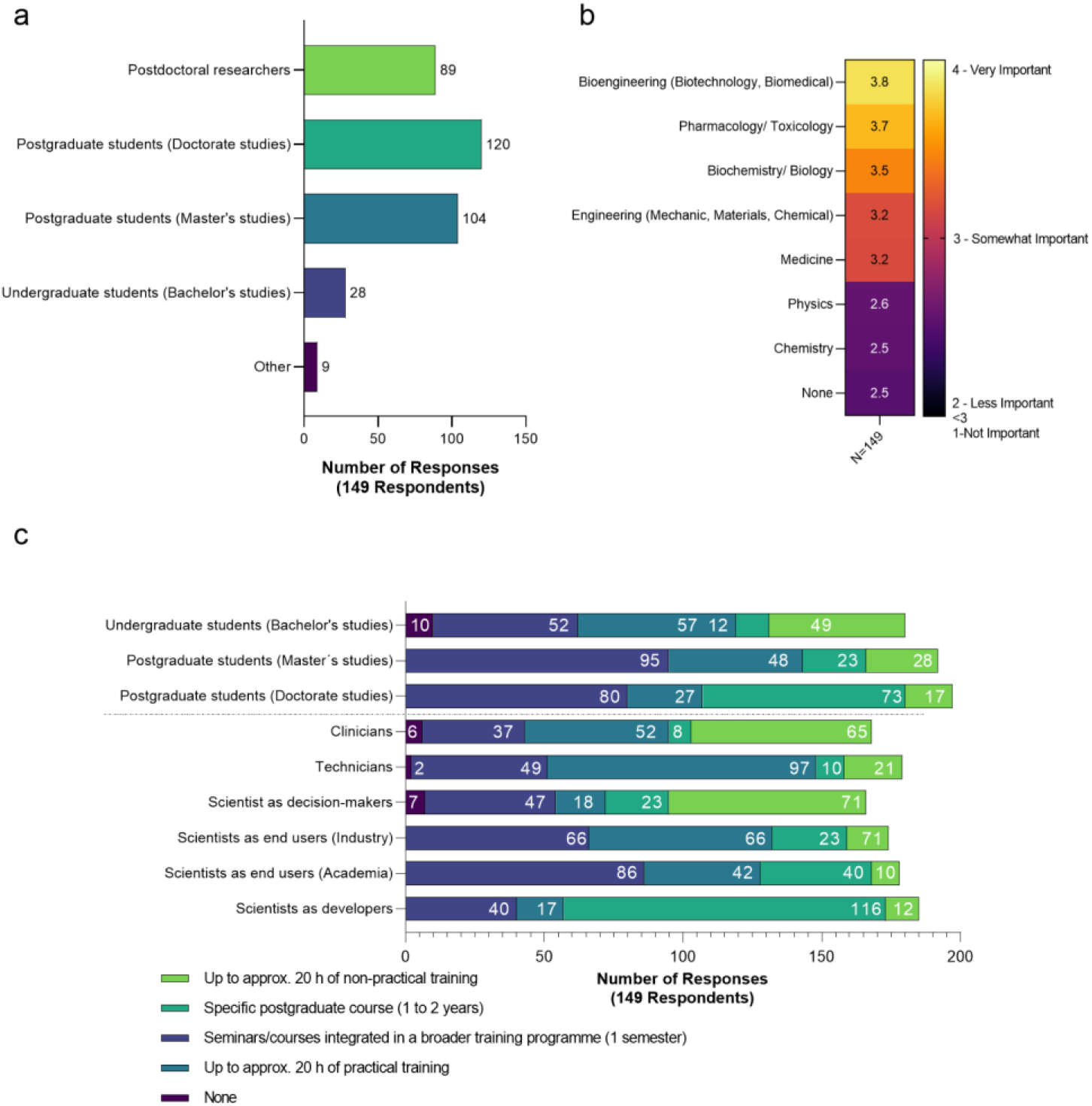
The extent of training in OoC technology to be given to various stakeholders: (a) at what educational level the stakeholder should receive training, (b) STEM courses where OoC topic could be integrated. The importance level was ranked by assigning values to the options to generate a ranking as following Very important = 4; Somewhat important = 3; Less important = 2; Not important = 1. We excluded respondents who skipped the question and those who answered “not sure”. The heatmap contains the average ranking per response and per group. (c) amount of theoretical and practical training each stakeholder should receive. The number of respondents for each specific amount of training is presented cumulatively inside of the respective bar. The data reflects the responses to Q6 (a), Q15 (b) and Q14 (c).

Respondents suggested *up to 20 hours of practical or non-practical training* at the bachelor’s level while at least *one semester* at the PhD level.

Lastly, to further analyze the expectations of the respondents towards training efforts, we sought to understand the respondents themselves, by analyzing their own career trajectory.

While most respondents held a bachelor’s degree in basic sciences, such as *biochemistry/biology* (38%) or *engineering* (21%), we observed a shift towards multidisciplinary fields for postgraduate studies (**Fig.5**). A majority of the *PhD candidates* held a bachelor’s, master’s (33%,46%) and were pursuing a PhD in *bioengineering. Senior scientists* on the other hand, held most commonly a bachelor’s and master’s in *biochemistry/biology* (46% and 31% respectively) and a doctorate degree in *bioengineering* (26%). The respondents in the *NAI category mostly held all three* degrees in *biochemistry/biology* (45% for bachelor’s, 32% for master’s and 23% for doctorate degrees). Researchers with *interdisciplinary expertise* gained it very early with a bachelor’s in *bioengineering* (35%), followed by graduate or postgraduate degrees in *bioengineering* as well (54 and 55% respectively). While fewer respondents with *engineering expertise* (74%) pursued a doctorate degree than those with *biology* or *interdisciplinary expertise* (88%, 92% respectively), they tended to switch towards bioengineering degrees at the masters and doctorate levels. Respondents with biology expertise spread across different disciplines such as *bioengineering, biochemistry/biology, pharmacology/toxicology* or *medicine* during doctorate studies. Moreover, 55% of them declared no familiarity with *microfabrication techniques* (**Fig.6 a**) while the group with engineering expertise noted reasonable familiarity with *biological systems* (**Fig.6 b**).

**Figure 5:**
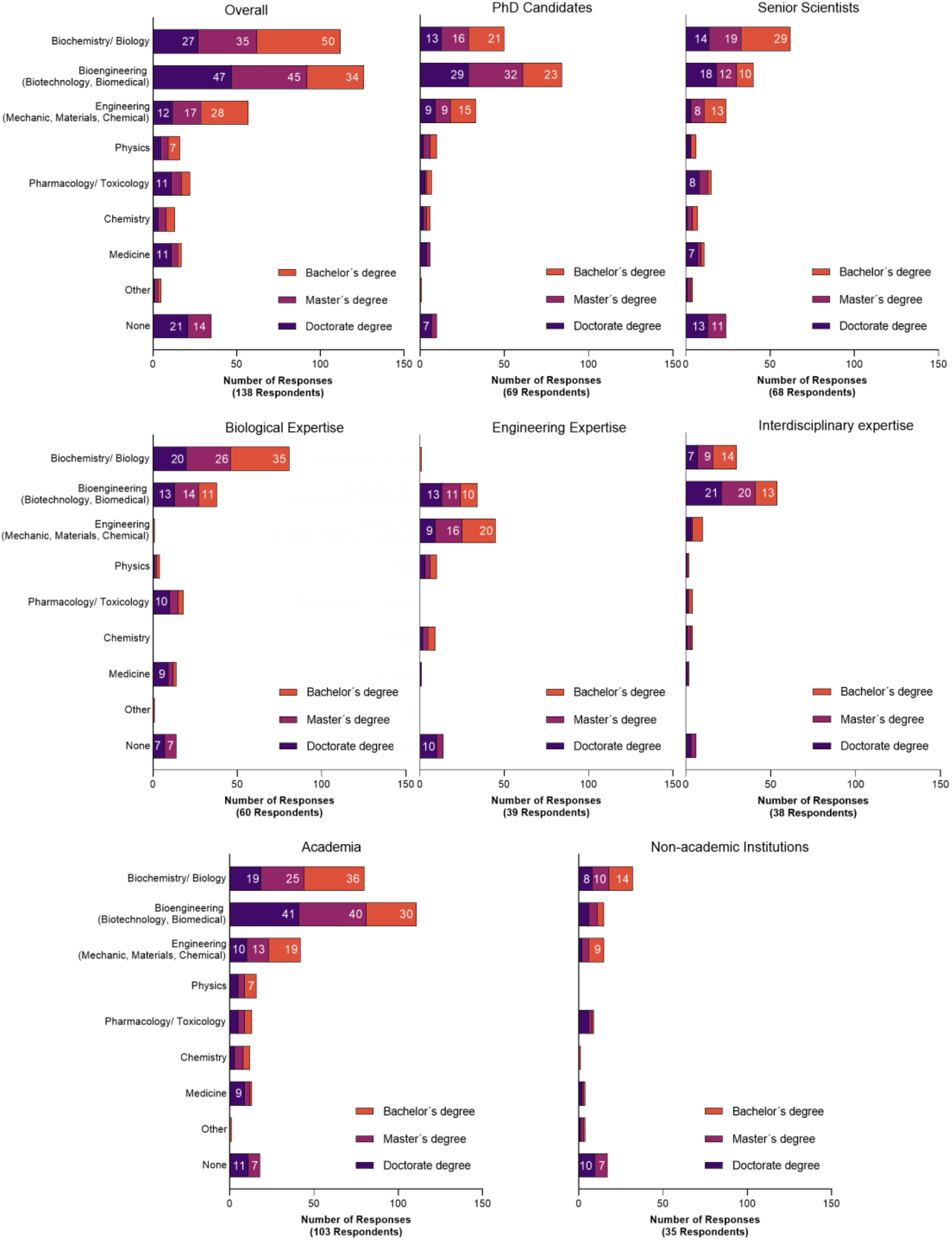
The academic background of all respondents (overall dataset) and the subgroups of *PhD candidates, senior scientists, academia, Non-academic institutions* and *biology/engineering/interdisciplinary experts*. The data reflects the responses to Q1.d e and f.

**Figure 6.**
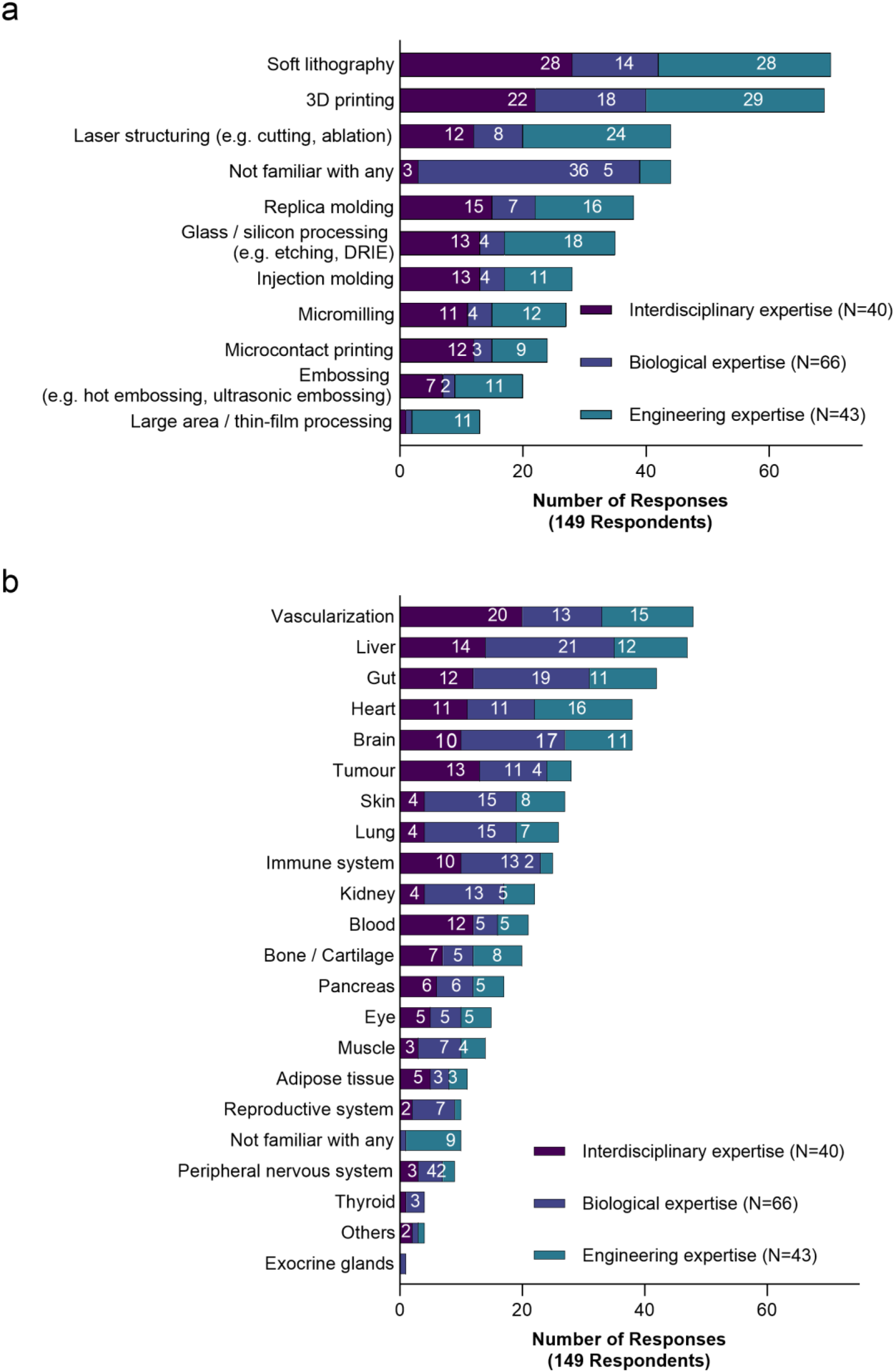
Familiarity of Interdisciplinary (N=40), Biological (N=66) and Engineering experts (N=43) in (a) microfabrication techniques (b) with biological systems. The number of respondents for each specific amount of training is presented cumulatively inside of the respective bars. The data reflects the responses to Q2 and Q3.

## Discussion

The overall answers to the questionnaire clearly showed that there is a strong need to train the next generation of OoC researchers. The key needs are to increase end-user familiarity with chip development and production and to train *technicians* and *scientists as end-users* in quality assurance and qualification of OoC models. Early-career researchers should be trained early on in OoC technology; courses should be included in applied science studies such as bioengineering, pharmacology/toxicology and biochemistry/biology.

The professional background of the respondents was essential to understand their answers and needs, and to draw conclusions about the direction that training in OoC technology should take for the next generation of researchers. Respondents currently enrolled in a doctoral program and actively involved in OoC research provided an important perspective that could guide training efforts given their current experiences. In our opinion, a young field such as OoC tends to have a larger presence in academia and it could be useful to include more insight from researchers in industry for topics in this field in future surveys. Due to the different research approaches of academia and NAI, we placed particular focus on the comparison of these respondent groups, which revealed different levels of importance of the individual training aspects and in terms of who should receive training. The relevance of interdisciplinarity was also addressed and importantly, the new generations of researchers are already obtaining interdisciplinary training earlier in their careers. This is probably not specific to the OoC field but essential to identify training needs and justify our analysis according to the respondents biological, engineering, or interdisciplinary expertise. We also profiled the respondents based on the type of research they work on as *basic, translational, and applied* research but we did not include the definitions of each research type. In an interdisciplinary field, those definitions may substantially vary between the biology experts and engineering experts. Therefore, a comparison between those groups was not performed. It would have been also of interest to stratify the respondents according to the OoC technologies development stage (platform development, OoC design, biological applications, or regulatory use) but that information was not available.

Beyond *training*, the aspects *usability, qualification* and *definition of cell culture standards* in the OoC field were deemed to be of high importance. Interestingly, the data also suggests that the respondents do not see the *comparison with animal* and *clinical data* as important aspects by ranking them as the least important aspects. Many efforts have been made recently towards qualification and standardization of OoC technologies with initiatives from academia, regulatory agencies and industry. A recent technical report published by the European Centre for the Validation of Alternative Methods (ECVAM) highlights the interest of the research community in establishing the scientific validity of complex *in vitro* models, particularly OoCs (Batista Leite et al., 2021). This report also stated that OoCs further require standardization at both technical and biological levels and that investment in training in aspects of both regulatory and biomedical research is essential. More recently, initiatives on the standardization of OoC have been taking place (Mastrangeli & van den Eijnden-van Raaij, 2021; Piergiovanni et al., 2021) such as the European consortium Moore4medical.eu developing open technology platforms (Mastrangeli et al., 2021). The WelcomeLeap foundation also launched the Hope program aiming at funding the development of organ platforms and connections. From the industry perspective, the Innovation & Quality Microphysiological Systems (IQ MPS) Affiliate, a consortium of pharmaceutical and biotechnology companies within the International Consortium for Innovation & Quality in Pharmaceutical Development (the IQ Consortium) identifies qualification as a key priority. Importantly, the IQ MPS states that qualification of OoCs, rather than validation in the OECD sense, is not trivial and cannot be dissociated from the context of use (Baran et al., 2022).

In the 3Rs context, a 1:1 replacement of animal models is no longer a priority and it is generally accepted that the key lies in developing models with human relevance, thus enhancing their translational value and their acceptance in the larger scientific community (Herrmann et al., 2019; Loskill et al., 2021). Possibly, the low importance attributed to *comparison with animal* and *clinical data* reflects this vision that microphysiological systems mimicking the complexity of the human body are able to generate knowledge with a mechanistic understanding and that using relevant controls and a scientific agreement of standards is more important than direct comparisons of significantly different model systems. The importance of using *in silico* modeling was also identified as somewhat important for the development of the OoC field (rank of 4.2 for *PKPD modeling)* and as an important training need for end users in industry with a rank 4.4-4.7 from respondents of all groups. Interestingly, almost half of the respondents (67/149), especially *PhD Candidates* and *engineering experts* responded “not sure” about the importance of *PKPD modeling* for training. *PKPD modeling* allows for correlation of *in vitro* data to *in vivo* responses (Cirit & Stokes, 2018; Maass et al., 2017) and possibly, this definition is not yet widely understood. A limitation of the employed questionnaire is the hiatus of questions with regards to the context of use of the OoC technologies, which could be a topic for future investigation. The context of use can shape the specific needs, which are possibly very different, for instance, for herbal products of food supplements industry, mainly interested in PKPD and toxicological data, compared to pharmaceutical industry, with a higher interest in extracting safety and efficacy mechanistic information from OoC models.

Overall, the respondents mainly worked in *applied* (106/149) and *basic research* (67/149) as well as the *pharmaceutical industry* (23/149) while other potential users in the *chemical and cosmetics industry* were underrepresented; potentially providing an explanation for the lower importance attributed to including training in OoC in chemistry study courses. This underrepresentation of industrial end users other than from pharmaceutical industry is mirrored in the overall OoC community, with the hesitancy and challenges of using novel complex model systems for regulatory toxicology being a possible reason.

Most of the respondents were expected to have experienced the challenges of working in multidisciplinary teams throughout their careers. Hence, interdisciplinary training was considered important for *scientists as developers* that connects the engineering aspects of the devices to their biological application. Moreover, training should start at early stages and should be integrated into interdisciplinary courses (*e.g. bioengineering*), to allow for sustainable development of this field. To the best of our knowledge, training activities in the bachelor’s and master’s levels of study were mostly unstructured and often limited to an introductory lecture integrated within a broader semester topics for different programs. Hands-on training for OoCs has also been provided directly by companies that sell microfluidic cell culture systems. Although hands-on training promoted by companies is a valuable contribution, it does not provide broad training and is often coupled with marketing interests. Such training actions are also commonly limited to a maximum of two-day sessions. PhD Candidates and postdoctoral researchers are the stakeholders with more diverse training opportunities. At a European level, training provided by the Netherlands Organ-on-Chip Initiative (NOCI) and the Marie Skłodowska-Curie Innovative Training Network (MSCA-ITN) EUROoC are good examples of strategies to gather diverse knowledge for training PhD candidates. Both initiatives bring together experts from stem cell biology, microfabrication, medical ethics, genetics, electronics, microfluidics and other fields to provide training for PhD candidates. These networks include partners from the academic sector, industrial sector, SMEs and regulatory authorities highlighting the importance of integrative training for the next generation of scientists.

Multidisciplinary, interdisciplinary, and transdisciplinary curriculum development and implementation is a recognized challenge (Budwig & Alexander, 2020; Takeuchi et al., 2020). Miranda *et al*. proposed to integrate four core components for a multidisciplinary training (Miranda et al., 2021): i) competencies (hard and soft skills), ii) learning methods (challenge- and problem-based learning), iii) information and communication technologies (digital tools and platforms), and iv) infrastructure (hands-on and class-room training). To approach interdisciplinary training at the institutional/university level, several concepts have been introduced, e.g., the creation of academic units within universities, so called *interdisciplinary organizations*, in parallel with the subject specific structure of departments (Yang et al., 2021). Interdisciplinary organizations allow to unify different STEM research areas (e.g., engineers, biochemists, toxicologists) to target a specific problem. Conversely, Takeuchi et al. suggest that STEM education should be transdisciplinary, defining it as more than the sum of the independent STEM disciplines and as an iterative and synergistic use of competencies from those disciplines (Takeuchi et al., 2020).

Respondents at the level of *PhD candidates* started an interdisciplinary career pathway earlier than *senior scientists*. Younger generations of scientists have likely recognized the importance of multidisciplinary training to achieve success in bridging biology and engineering concepts to develop or apply OoC technologies. However, the academic career choices of *senior scientists* working in the OoC field differed, suggesting that most of the *senior scientists* started engaging in multidisciplinary training later in their career and probably not through a formal academic degree. *PhD candidates* also considered it more important to promote awareness and uptake of by scientists from other fields.

The higher interdisciplinary background of *Academia* compared to *NAI* may be the reason why the responses of *academia* leaned towards engineering topics and promoting the uptake by other scientists while *NAI* considered that training in cell culture and translational topics were needed over engineering. This suggests that *scientists as developers* likely worked in academia while *NAIs* mainly comprise of *endusers* of OoC products. Respondents in *NAIs* has a deeper *pharmacology/toxicology* background while *academia* has a *bioengineering* background – two important aspects for OoC application. This points towards an opportunity where training efforts could bridge these currently separate sets of expertise through the sharing of knowledge (Esch et al., 2015; Holley et al., 2016).

Respondents with *interdisciplinary expertise* attributed higher importance to training in both biological and engineering aspects. Another striking difference is that *engineering experts* showed a certain degree of familiarity with biological systems that is essential for proper OoC design and fabrication but are not sure or did not respond concerning the importance of *PKPD modeling* (27/39). Contrarily, the understanding of the microfabrication techniques and bioengineering principles seemed not to be essential for *biological experts*, who likely considered themselves as *end users*. However, a better understanding of the material, design and fabrication concepts and limitations from the end users could bring a faster and more productive development of innovative OoC. These observations could point towards a potential training avenue targeting the biology experts. Importantly, the OoC field is an example of the emerging need for transdisciplinary education (J. Beebe et al., 2013; Ramadan & Zourob, 2020). To be an OoC expert one needs biological and engineering expertise and needs to be able to see though both lenses in an integrated way and using a common language (Mastrangeli & van den Eijnden-van Raaij, 2021).

Based on a thorough analysis of responses from more than 100 experts with different scientific backgrounds and from different sectors, we drew several key conclusions and provide an answer to the three overarching questions formulated initially:

1. Which **aspects of training** are the most important for the overall development of the OoC field?

a. *Definition of cell culture standards*
b. OoC *usability*
2. Which **specific type of training** in the OoC field should each stakeholder receive?

a. *Scientists as end users* with a biology background should be trained in *microfabrication and microfluidics principles*
b. *Scientists as developers* should be trained in *pharmacology/toxicology*
3. What is the **amount of training** that each stakeholder should receive to achieve a certain level of experience in the OoC field?

a. *Scientists as developers* should *undergo specific post graduate training (1-2 years)*
b. *Scientists as end users should* undergo specific *practical training (up to 20h)* and *seminars/courses integrated in a broader training*

The information collected and analyzed in this study could foster STEM undergraduate and graduate education in this technology and result in well-qualified and competent transdisciplinary professionals, who will continue exploring and successfully applying OoC technology.

## Acknowledgements

The authors kindly thank all the respondents for their time and for their help in distributing the questionnaire with a special thanks to Janny van den Eijnden-van Raaij for her advice and support during the drafting of the questionnaire.

## List of Abbreviations

3Rs: Replacement, Reduction and Refinement
ECVAM: European Centre for the Validation of Alternative Methods
IQ MPS: Innovation & Quality Microphysiological Systems
OECD: Organisation for Economic Co-operation and Development
OoC: Organ-on-chip
PKPD: pharmacokinetic/pharmacodynamic
R&D: Research and Development
SME: Small and Medium Enterprise
STEM: Science, Technology, Engineering, and Mathematics

## Data availability

All data and analysis code is available in the supplementary files

