## supplementary file 1 for "Training the next generation of researchers in the Organ-on-Chip field"

### Questionnaire for stakeholders: Training needs of the next generation of researchers and technicians in Organ-on-a-Chip field

Fields marked with * are mandatory.

#### Questionnaire for stakeholders

#### Organ-on-a-chip

#### Training needs of the next generation of researchers and technicians

This questionnaire aims at surveying the **training needs** of the Organ-on-Chip community to **promote the Organ-on-Chip systems qualification, usability, uptake and long-term development** in a variety of fields.
Organ-on-Chip applications in Basic Research, Pharmaceutical Drug Development, Safety assessment of Drugs, Cosmetics and Chemicals among others are facing an exponential arise in interest. Therefore, **speci fic training is required** on the production of such cell culture systems using advanced microfabrication techniques and adequate on-chip characterization of relevant cell functions. This survey is directed to those we considered to be the current and future strategic stakeholders in the advance and use of Organ-on-Chip. On the one hand, we aim at **preparing scientists and technicians for new types of employment** that will arise while, on the other hand, **providing industry and academia with professionals able to keep up with innovation** in the field. The answers to this survey will contribute to designing appropriate training programs to fulfil the needs of this emerging field.
The data resulting from the survey will be publicly available on the **ORCHID website** and disseminated on the project **twitter** account.
This survey contais 15 questions and will take around **15 minutes** of your time.

The privacy statement, available below, outlines in detail how the data that you provide as part of this survey will be protected.

Privacy_Statement.pdf

☐ I agree with the privacy statement.

#### Section 1 – Professional profile

* 1. How would you **define yourself as a professional**?

a) **Type of institution**

*at most 2 choice(s)*

- [ ] Academia
- [ ] Industry
- [ ] SME (as defined by H2020)
- [ ] Governamental Organization
- [ ] Non-Governamental Organization
- [ ] Hospitals
- [ ] Other (please specify)

Other:

* b) **Job Level**

*at most 1 choice(s)*

- [ ] Head of Institution/Top Management
- [ ] Senior Officer/Middle Management/Principal Investigator/Associate Professor
- [ ] Technical Officer/Staff Scientist/Post-doctoral Fellow
- [ ] PhD Student / Doctoral Candidate
- [ ] Other (please specify) ________________________________

Other:

* c) **Field of work**

*at most 2 choice(s)*

- [ ] Basic Research
- [ ] Applied Research
- [ ] Pharmaceutical Industry
- [ ] Clinic Cosmetics or Chemical Industry
- [ ] Food Industry
- [ ] Other (please specify) ________________________________

Other:

* d) **Field of Graduate University education** (Bachelor's Degree)

*at most 1 choice(s)*

- [ ] Engineering (Mechanic, Materials, Chemical)
- [ ] Bioengineering (Biotechnology, Biomedical)
- [ ] Physics
- [ ] Chemistry
- [ ] Biochemistry/ Biology
- [ ] Medicine

- [ ] Pharmacology/ Toxicology
- [ ] None
- [ ] Other (please specify) ________________________________

Other:

* e) **Field of first Postgraduate University education** (Master's Degree)

*at most 1 choice(s)*

- [ ] Engineering (Mechanic, Materials, Chemical)
- [ ] Bioengineering (Biotechnology, Biomedical)
- [ ] Physics
- [ ] Chemistry
- [ ] Biochemistry/ Biology
- [ ] Medicine
- [ ] Pharmacology/ Toxicology
- [ ] None
- [ ] Other (please specify) ________________________________

Other:

* f) **Field of second Postgraduate University education** (Doctorate Degree)

*at most 1 choice(s)*

- [ ] Engineering (Mechanic, Materials, Chemical)
- [ ] Bioengineering (Biotechnology, Biomedical)
- [ ] Physics
- [ ] Chemistry
- [ ] Biochemistry/ Biology
- [ ] Medicine
- [ ] Pharmacology/ Toxicology
- [ ] None
- [ ] Other (please specify) ________________________________

Other:

* g) **Main area of expertise**

*at most 2 choice(s)*

- [ ] Materials Science
- [ ] Microfabrication / Microfluidics
- [ ] Automation / Process engineering
- [ ] Cell Biology / Molecular Biology

- [ ] Genomics / Multi-omics
- [ ] Pharmacology / Toxicology
- [ ] Clinical Sciences
- [ ] Other (please specify)_______________

Other:

* 2. Select the **tissues/organs/systems/biological functions** with which you are familiar and/or you work with? (Select all that apply)

| | | | |
|---|---|---|---|
| ☐ Skin | ☐ Brain | ☐ Pancreas | ☐ Reproductive System |
| ☐ Eye | ☐ Peripheral nervous system | ☐ Thyroid | ☐ Exocrine glands |
| ☐ Heart | ☐ Tumour | ☐ Muscle | ☐ Not familiar with any specific tissues/organs/systems /biological functions |
| ☐ Liver | ☐ Vascularization | ☐ Kidney | ☐ Others |
| ☐ Gut | ☐ Immune System | ☐ Lung | |
| ☐ Adipose tissue | ☐ Blood | ☐ Bone/Cartilage | |

* 3. Select the **microfabrication techniques** for polymer-based microfluidic devices with which you are familiar and/or you work with?

- [ ] Soft Lithography
- [ ] Glass / silicon processing (e.g. etching, DRIE)
- [ ] Large area / thin-film processing
- [ ] Embossing (e.g. hot embossing, ultrasonic embossing)
- [ ] Replica molding
- [ ] Microcontact printing
- [ ] 3D printing
- [ ] Micromilling
- [ ] Laser structuring (e.g. cutting, ablation)
- [ ] Injection molding
- [ ] Not familiar with any microfabrication technique
- [ ] Others (please specify)_______

Other:

#### Section 2 – Opinion on the state of Organ-on-Chip field development

4. **How important** are the following aspects for the Organ-on-Chip field development?

*between 12 and 13 answered rows*

| | Very Important | Somewhat Important | Less Important | Not Important | Not Sure |
|---|---|---|---|---|---|
| Microfabrication techniques | ○ | ○ | ○ | ○ | ○ |
| Production scale-up of Organ-on-Chip systems | ○ | ○ | ○ | ○ | ○ |
| Definition of specific cell culture standards – function and origin of cells | ○ | ○ | ○ | ○ | ○ |
| Sensors integration and real-time monitoring | ○ | ○ | ○ | ○ | ○ |
| High throughput cultivation and endpoint measurements | ○ | ○ | ○ | ○ | ○ |
| PKPD modeling | ○ | ○ | ○ | ○ | ○ |
| Qualification of the models | ○ | ○ | ○ | ○ | ○ |
| Usability | ○ | ○ | ○ | ○ | ○ |
| Comparision with clinical data | ○ | ○ | ○ | ○ | ○ |
| Comparision with animal data | ○ | ○ | ○ | ○ | ○ |
| Uptake by scientists from other fields | ○ | ○ | ○ | ○ | ○ |
| Training | ○ | ○ | ○ | ○ | ○ |
| Other (please specify) | ○ | ○ | ○ | ○ | ○ |

Other:

#### Section 3 – Specific training needs

5. **How important** is it to provide specific training for each of the following stakeholders, to promote the Organ-on-Chip systems qualification, usability, uptake and/or long-term development?

*between 6 and 7 answered rows*

| | Very Important | Somewhat Important | Less Important | Not Important | Not Sure |
|---|---|---|---|---|---|
| Scientists as developers | ○ | ○ | ○ | ○ | ○ |
| Scientists as end users (Academia) | ○ | ○ | ○ | ○ | ○ |
| Scientists as end users (Industry) | ○ | ○ | ○ | ○ | ○ |
| Scientists as decision-makers (Regulators, Grant evaluators or peer reviewers) | ○ | ○ | ○ | ○ | ○ |

| | | | | | |
|---|---|---|---|---|---|
| Technicians | ○ | ○ | ○ | ○ | ○ |
| Clinicians | ○ | ○ | ○ | ○ | ○ |
| Other (please specify) | ○ | ○ | ○ | ○ | ○ |

Other:

* 6. **At which level** do you consider that specific training is necessary to promote the Organ-on-Chip systems qualification, usability, uptake and/or long-term development?

- ☐ Postdoctoral researchers
- ☐ Postgraduate Students (Doctorate studies)
- ☐ Postgraduate Students (Master's studies)
- ☐ Undergraduate Students (Bachelor's Studies)
- ☐ Other (please specify)______________

Other:

7. **How important** are the following elements to consider for training, for **Scientists as developers**, regarding the improvement of Organ-on-Chip systems qualification, usability, uptake and/or long-term development?

*between 13 and 14 answered rows*

| | Very Important | Somewhat Important | Less Important | Not Important | Not Sure |
|---|---|---|---|---|---|
| Biomaterials | ○ | ○ | ○ | ○ | ○ |
| Microfabrication techniques and manufacturability | ○ | ○ | ○ | ○ | ○ |
| Microfluidic principles | ○ | ○ | ○ | ○ | ○ |
| Cell culture and stem cell technology | ○ | ○ | ○ | ○ | ○ |
| Bio banking, Data Management and Protection | ○ | ○ | ○ | ○ | ○ |
| Monitoring and analysing (molecular biology / omics) | ○ | ○ | ○ | ○ | ○ |
| Monitoring and analysing (sensors, imaging) | ○ | ○ | ○ | ○ | ○ |
| PKPD modelling | ○ | ○ | ○ | ○ | ○ |
| Pharmacology and Toxicology principles | ○ | ○ | ○ | ○ | ○ |

| | | | | | |
|---|---|---|---|---|---|
| Quality Assurance | ◯ | ◯ | ◯ | ◯ | ◯ |
| Science Communication | ◯ | ◯ | ◯ | ◯ | ◯ |
| Regulatory affairs | ◯ | ◯ | ◯ | ◯ | ◯ |
| Ethics | ◯ | ◯ | ◯ | ◯ | ◯ |
| Other (please specify) | ◯ | ◯ | ◯ | ◯ | ◯ |

Other:

8. **How important** are the following elements to consider for training, for **Scientists as end users (Academia)**, regarding the improvement of Organ-on-Chip systems qualification, usability, uptake and/or long-term development?

*between 13 and 14 answered rows*

| | Very Important | Somewhat Important | Less Important | Not Important | Not Sure |
|---|---|---|---|---|---|
| Biomaterials | ◯ | ◯ | ◯ | ◯ | ◯ |
| Microfabrication techniques and manufacturability | ◯ | ◯ | ◯ | ◯ | ◯ |
| Microfluidic principles | ◯ | ◯ | ◯ | ◯ | ◯ |
| Cell culture and stem cell technology | ◯ | ◯ | ◯ | ◯ | ◯ |
| Bio banking, Data Management and Protection | ◯ | ◯ | ◯ | ◯ | ◯ |
| Monitoring and analysing (molecular biology / omics) | ◯ | ◯ | ◯ | ◯ | ◯ |
| Monitoring and analysing (sensors, imaging) | ◯ | ◯ | ◯ | ◯ | ◯ |
| PKPD modelling | ◯ | ◯ | ◯ | ◯ | ◯ |
| Pharmacology and Toxicology principles | ◯ | ◯ | ◯ | ◯ | ◯ |
| Quality Assurance | ◯ | ◯ | ◯ | ◯ | ◯ |
| Science Communication | ◯ | ◯ | ◯ | ◯ | ◯ |
| Regulatory affairs | ◯ | ◯ | ◯ | ◯ | ◯ |
| Ethics | ◯ | ◯ | ◯ | ◯ | ◯ |
| Other (please specify) | ◯ | ◯ | ◯ | ◯ | ◯ |

Other:

9. **How important** are the following elements to consider for training, for **Scientists as end users (Industry)**, regarding the improvement of Organ-on-Chip systems qualification, usability, uptake and/or long-term development?

*between 13 and 14 answered rows*

| | Very Important | Somewhat Important | Less Important | Not Important | Not Sure |
|---|---|---|---|---|---|
| Biomaterials | ○ | ○ | ○ | ○ | ○ |
| Microfabrication techniques and manufacturability | ○ | ○ | ○ | ○ | ○ |
| Microfluidic principles | ○ | ○ | ○ | ○ | ○ |
| Cell culture and stem cell technology | ○ | ○ | ○ | ○ | ○ |
| Bio banking, Data Management and Protection | ○ | ○ | ○ | ○ | ○ |
| Monitoring and analysing (molecular biology / omics) | ○ | ○ | ○ | ○ | ○ |
| Monitoring and analysing (sensors, imaging) | ○ | ○ | ○ | ○ | ○ |
| PKPD modelling | ○ | ○ | ○ | ○ | ○ |
| Pharmacology and Toxicology principles | ○ | ○ | ○ | ○ | ○ |
| Quality Assurance | ○ | ○ | ○ | ○ | ○ |
| Science Communication | ○ | ○ | ○ | ○ | ○ |
| Regulatory affairs | ○ | ○ | ○ | ○ | ○ |
| Ethics | ○ | ○ | ○ | ○ | ○ |
| Other (please specify) | ○ | ○ | ○ | ○ | ○ |

Other:

10. **How important** are the following elements to consider for training, for **Scientists as decision-makers (Regulators, Grant evaluators or peer reviewers)**, regarding the improvement of Organ-on-Chip systems qualification, usability, uptake and/or long-term development?

*between 13 and 14 answered rows*

| | Very Important | Somewhat Important | Less Important | Not Important | Not Sure |
|---|---|---|---|---|---|

| | | | | | |
|---|---|---|---|---|---|
| Biomaterials | ○ | ○ | ○ | ○ | ○ |
| Microfabrication techniques and manufacturability | ○ | ○ | ○ | ○ | ○ |
| Microfluidic principles | ○ | ○ | ○ | ○ | ○ |
| Cell culture and stem cell technology | ○ | ○ | ○ | ○ | ○ |
| Bio banking, Data Management and Protection | ○ | ○ | ○ | ○ | ○ |
| Monitoring and analysing (molecular biology / omics) | ○ | ○ | ○ | ○ | ○ |
| Monitoring and analysing (sensors, imaging) | ○ | ○ | ○ | ○ | ○ |
| PKPD modelling | ○ | ○ | ○ | ○ | ○ |
| Pharmacology and Toxicology principles | ○ | ○ | ○ | ○ | ○ |
| Quality Assurance | ○ | ○ | ○ | ○ | ○ |
| Science Communication | ○ | ○ | ○ | ○ | ○ |
| Regulatory affairs | ○ | ○ | ○ | ○ | ○ |
| Ethics | ○ | ○ | ○ | ○ | ○ |
| Other (please specify) | ○ | ○ | ○ | ○ | ○ |

Other:

11. **How important** are the following elements to consider for training, for **Technicians**, regarding the improvement of Organ-on-Chip systems qualification, usability, uptake and/or long-term development?

*between 13 and 14 answered rows*

| | Very Important | Somewhat Important | Less Important | Not Important | Not Sure |
|---|---|---|---|---|---|
| Biomaterials | ○ | ○ | ○ | ○ | ○ |
| Microfabrication techniques and manufacturability | ○ | ○ | ○ | ○ | ○ |
| Microfluidic principles | ○ | ○ | ○ | ○ | ○ |
| Cell culture and stem cell technology | ○ | ○ | ○ | ○ | ○ |
| Bio banking, Data Management and Protection | ○ | ○ | ○ | ○ | ○ |

| | | | | | |
|---|---|---|---|---|---|
| Monitoring and analysing (molecular biology / omics) | ○ | ○ | ○ | ○ | ○ |
| Monitoring and analysing (sensors, imaging) | ○ | ○ | ○ | ○ | ○ |
| PKPD modelling | ○ | ○ | ○ | ○ | ○ |
| Pharmacology and Toxicology principles | ○ | ○ | ○ | ○ | ○ |
| Quality Assurance | ○ | ○ | ○ | ○ | ○ |
| Science Communication | ○ | ○ | ○ | ○ | ○ |
| Regulatory affairs | ○ | ○ | ○ | ○ | ○ |
| Ethics | ○ | ○ | ○ | ○ | ○ |
| Other (please specify) | ○ | ○ | ○ | ○ | ○ |

Other:

12. **How important** are the following elements to consider for training, for **Clinicians**, regarding the improvement of Organ-on-Chip systems qualification, usability, uptake and/or long-term development?

*between 13 and 14 answered rows*

| | Very Important | Somewhat Important | Less Important | Not Important | Not Sure |
|---|---|---|---|---|---|
| Biomaterials | ○ | ○ | ○ | ○ | ○ |
| Microfabrication techniques and manufacturability | ○ | ○ | ○ | ○ | ○ |
| Microfluidic principles | ○ | ○ | ○ | ○ | ○ |
| Cell culture and stem cell technology | ○ | ○ | ○ | ○ | ○ |
| Bio banking, Data Management and Protection | ○ | ○ | ○ | ○ | ○ |
| Monitoring and analysing (molecular biology / omics) | ○ | ○ | ○ | ○ | ○ |
| Monitoring and analysing (sensors, imaging) | ○ | ○ | ○ | ○ | ○ |
| PKPD modelling | ○ | ○ | ○ | ○ | ○ |
| Pharmacology and Toxicology principles | ○ | ○ | ○ | ○ | ○ |
| Quality Assurance | ○ | ○ | ○ | ○ | ○ |

| | | | | | |
|---|---|---|---|---|---|
| Science Communication | ○ | ○ | ○ | ○ | ○ |
| Regulatory affairs | ○ | ○ | ○ | ○ | ○ |
| Ethics | ○ | ○ | ○ | ○ | ○ |
| Other (please specify) | ○ | ○ | ○ | ○ | ○ |

Other:

13. What is the most adequate **complexity level of specific training** for each of the following stakeholders, to promote Organ-on-Chip systems qualification, usability, uptake and/or long-term development? (Select the most adequate)

*between 6 and 7 answered rows*

| | Deep knowledge /Theoretical and practical skills | Competence/ Theoretical skills | Competence /Practical skills | Introductory/ Awareness | None |
|---|---|---|---|---|---|
| Scientists as developers | ☐ | ☐ | ☐ | ☐ | ☐ |
| Scientists as end users (Academia) | ☐ | ☐ | ☐ | ☐ | ☐ |
| Scientists as end users (Industry) | ☐ | ☐ | ☐ | ☐ | ☐ |
| Scientists as decision-makers (Regulators, Grant evaluators or peer reviewers) | ☐ | ☐ | ☐ | ☐ | ☐ |
| Technicians | ☐ | ☐ | ☐ | ☐ | ☐ |
| Clinicians | ☐ | ☐ | ☐ | ☐ | ☐ |
| Other (please specify) | ☐ | ☐ | ☐ | ☐ | ☐ |

Other:

14. What is the most adequate **amount of specific training** for each of the following stakeholders, to promote Organ-on-Chip systems qualification, usability, uptake and/or long-term development? (Select the most adequate)

*between 9 and 10 answered rows*

| | Specific postgraduate course (1 to 2 years) | Seminars/courses integrated in a broader training programme (1 semester) | Up to approx. 20 h of practical training | Up to approx. 20 h of non-practical training | None |
|---|---|---|---|---|---|
| Scientists as developers | ☐ | ☐ | ☐ | ☐ | ☐ |
| Scientists as end users (Academia) | ☐ | ☐ | ☐ | ☐ | ☐ |
| Scientists as end users (Industry) | ☐ | ☐ | ☐ | ☐ | ☐ |
| Scientists as decision-makers (Regulators, Grant evaluators or peer reviewers) | ☐ | ☐ | ☐ | ☐ | ☐ |
| Technicians | ☐ | ☐ | ☐ | ☐ | ☐ |
| Clinicians | ☐ | ☐ | ☐ | ☐ | ☐ |
| Postgraduate Students (Doctorate studies) | ☐ | ☐ | ☐ | ☐ | ☐ |
| Postgraduate Students (Master's studies) | ☐ | ☐ | ☐ | ☐ | ☐ |

| | | | | | |
|---|---|---|---|---|---|
| Undergraduate Students (Bachelor's Studies) | ☐ | ☐ | ☐ | ☐ | ☐ |
| Other (please specify) | ☐ | ☐ | ☐ | ☐ | ☐ |

Other:

15. **How important** would it be to include the topic of Organ-on-Chip technologies as a **seminar of course in the following broader field(s) of education**:

*between 7 and 8 answered rows*

| | Very Important | Somewhat Important | Less Important | Not Important | Not Sure |
|---|---|---|---|---|---|
| Engineering (Mechanic, Materials, Chemical) | ○ | ○ | ○ | ○ | ○ |
| Bioengineering (Biotechnology, Biomedical) | ○ | ○ | ○ | ○ | ○ |
| Physics | ○ | ○ | ○ | ○ | ○ |
| Chemistry | ○ | ○ | ○ | ○ | ○ |
| Biochemistry/ Biology | ○ | ○ | ○ | ○ | ○ |
| Medicine | ○ | ○ | ○ | ○ | ○ |
| Pharmacology/ Toxicology | ○ | ○ | ○ | ○ | ○ |
| None | ○ | ○ | ○ | ○ | ○ |
| Other (please specify) ______________________________ | ○ | ○ | ○ | ○ | ○ |

Other:

If you consider other relevant topics not covered by this questionnaire, **please leave you comments here**.
Choose to leave your comments anonymus or add your Name and Contact details.

If you have questions regarding this questionnaire, please contact
